# Genomic diversity and molecular epidemiology of a multidrug resistant *Pseudomonas aeruginosa* DMC30b isolated from hospitalized burn patient in Bangladesh

**DOI:** 10.1101/2022.07.06.498939

**Authors:** M. Nazmul Hoque, M. Ishrat Jahan, M. Anwar Hossain, Munawar Sultana

## Abstract

**Objectives:** *Pseudomonas aeruginosa* is a key opportunistic pathogen causing a wide range of community- and hospital-acquired infections in immunocompromised or catheterized patients. Here, we report the complete genome sequence of a multidrug resistant (MDR) *P. aeruginosa* DMC30b in order to elucidate the genetic diversity, molecular epidemiology, and underlying mechanisms for antimicrobial resistance and virulence.

**Methods:** *P. aeruginosa* DMC30b was isolated from septic wound swab of a severe burn patient. Whole-genome sequencing (WGS) was performed under Ion Torrent platform. The genome was annotated using the SPAdes v. 3.12.01 in an integrated Genome Analysis Platform (IonGAP) for Ion Torrent sequence data. The genome was annotated using the NCBI Prokaryotic Genome Annotation Pipeline (PGAP). *In-silico* predictions of antimicrobial resistance genes (ARGs), virulence factor genes (VFGs) and metabolic functional potentials were performed using different curated bioinformatics tools.

**Results:** *P. aeruginosa* DMC30b was classified as MDR and belongs to sequence type 244 (ST244). The complete genome size is 6,994,756 bp with a coverage of 76.76x, G+C content of 65.7% and a BUSCO (Benchmarking Universal Single-Copy Orthologs) score of 100. The genome of *P. aeruginosa* DMC30b harboured two plasmids (e,g., IncP-6 plasmid p10265-KPC; 78,007 bp and ColRNAI_pkOIISD1; 9,359 bp), 35 resistomes (ARGs) conferring resistance to 18 different antibiotics (including four beta-lactam classes), and 214 VFGs. It was identified as the 167^th^ ST244 strain among ∼ 5,800 whole-genome sequences of *P. aeruginosa* available in the NCBI database.

**Conclusion:** *P. aeruginosa* DMC30b belongs to ST244 and was identified as the 167^th^ such isolate to be submitted to NCBI, and the first complete ST244 genome from Bangladesh. The complete genome data with high genetic diversity and underlying mechanisms for antimicrobial resistance and virulence of *P. aeruginosa* DMC30b (ST244) will aid in understanding the evolution and phylogeny of such high-risk clones and provide a solid basis for further research on MDR or extensively drug resistant strains.

## 1. Background

*Pseudomonas aeruginosa* is an opportunistic Gram-negative, non-fermenting bacterial pathogen that causes a wide array of life-threatening, acute and chronic infections in hospitalized and immunocompromised patients, especially in the intensive care unit (ICU) [1, 2]. It is well established that bacterial proliferation in wounds contributes to infection and delayed wound healing [3]. The microenvironment of chronic wound is ideal for bioburden and usually contains multiple bacterial species [4, 5]. *P. aeruginosa* is one of the most common bacteria isolated from chronic wounds [6], and is highly prevalent in chronic wounds, estimated to be present in about 25% of all cases [7]. Infections caused by *P. aeruginosa* are usually difficult to treat and persistent due to the characteristic high frequency of emergence of MDR or extensively drug-resistant (XDR) strains [8-10]. The World Health Organization and Centers for Disease Control and Prevention have both designated *P. aeruginosa* as one of the major (critical) pathogens for which new antibiotics are desperately needed [11, 12]. Moreover, *P. aeruginosa* is a persistent and difficult-to-treat pathogen in many patients, and possesses a versatile arsenal of antimicrobial resistance determinants and virulence factors that enable survival, adaptation, and consequent persistence within the complex milieu of infections [13]. The increasingly frequent infections caused by MDR and XDR strains with limited therapeutic options are associated with high morbidity and mortality worldwide [13, 14]. Particularly the emergence of MDR and XDR strains due to bacterial expression of resistance genes such as β-lactamases, 16S rRNA methylases, and carbapenemases in recent years leading to severe infections with serious global threats to human health, emphasizing the need for novel (antibiotics-independent) treatment strategies. The most troublesome acquired resistance of *P. aeruginosa* is the production of carbapenemases, which confer resistance to most commercially available β-lactams [9]. The presence of these carbapenemases in high-risk clones and sequence types (STs) on complete genome or plasmid or the chromosome of *P. aeruginosa*, may be the cause of its successful dissemination in Bangladesh and beyond. Recent molecular and genomic studies reported that most of the infectious and virulent clones of *P. aeruginosa* belonged to ST235, ST244, ST308, ST1006, and ST1060 [15-17].

Recently, an increasing prevalence of MDR and XDR *P. aeruginosa* strains, with rates of between 15% and 30% in some geographical areas have been reported [18, 19]. Microbial and host factors furthermore impact the ongoing adaptive response exhibited by MDR, XDR and mutating strains of *P. aeruginosa* in immunocompromised and hospitalized patients. Furthermore, acquisition of advantageous attributes such as virulence factor assembly, motility, antibiotic resistance, and metabolic adaptation of the strains are the major contributors to morbidity and mortality [13]. Several molecular and typing methods have been used to study the evolution and genetic heterogeneity of *P. aeruginosa* because it is characterized by high genetic diversity [15, 20, 21]. In recent years, whole genome sequencing (WGS) and RNAseq analysis has enabled the study of isolates collected sequentially from patients and the characterization of the molecular epidemiology and evolutionary patterns and/or trajectories that comprise the hallmark complexity of diverse subclones of *P. aeruginosa* in clinically infected patients [15, 20, 22]. The study of molecular epidemiology and genetic diversity in MDR *P. aeruginosa* that harbour genes conferring resistance to a wide range of antibiotics and virulence factors, is key to understand the role in the dissemination of such resistance determinants among clinical and environmental isolates [2, 15]. The International *Pseudomonas aeruginosa* Consortium was formed with the aim of genome sequencing >1000 *P. aeruginosa* genomes and constructing an analysis pipeline for the study of *P. aeruginosa* evolution, virulence and antibiotic resistance [23]. Genomics data of this consortium will support molecular epidemiology for the surveillance of outbreaks and has the potential for future genotypic antimicrobial susceptibility testing as well as the identification of novel therapeutic targets and prognostic markers [24]. Herein this article, we describe the epidemiological distribution, host range, genomic diversities, ARGs and virulence factors found in an MDR *P. aeruginosa* DMC30b isolated from septic wound swab of a severely burn patient hospitalized in the Dhaka Medical College (DMC), Bangladesh. To elucidate the genetic diversity, molecular epidemiology, and underlying mechanisms for antimicrobial resistance and virulence of *P. aeruginosa* DMC30b, high throughput WGS and downstream bioinformatic analysis were performed.

## 2. Materials and Methods

### 2.1. Ethics statement

The study was approved by the Ethical Review Committee (ERC), Faculty of Biological Sciences, University of Dhaka, Bangladesh (reference 64/Biol.Scs./2018–2019) and carried out under the direct supervision of the laboratory biosafety officer. As the samples were collected from Dhaka Medical College Hospital, we did not handle the human subjects directly.

### 2.2. Isolate retrieval and antimicrobial susceptibility tests

*P. aeruginosa* DMC30b, previously isolated from septic wound swab of a severe burn patient [9], was retrieved and screened for antimicrobial susceptibility tests, genomic diversity and molecular epidemiological analysis. The isolate was plated onto the Mueller Hinton agar (MHA) (Oxoid, England), and incubated at 37 °C for 18-24 h followed by retrieval of pure colonies. The isolate was tested against a series of antibiotics recommended in the Clinical and Laboratory Standards Institute (CLSI) document M1007 [25] using the Kirby–Bauer method. Twenty antibiotics of 11 antibiotic groups were tested: imipenem (10 mg), meropenem (10 mg), doripenem (10 mg), chloramphenicol (30 mg), ampicillin (10 mg), doxycycline (30 mg), nitrofurantoin (300 mg), gentamicin (10 mg), trimethoprim (5 mg), tetracycline (30 mg), cefalexin (first generation; 30 mg), cefuroxime (second generation; 30 mg), cefotaxime (third generation; 30 mg), cefepime (fourth generation; 30 mg), nalidixic acid (first generation; 30 mg), ciprofloxacin (second generation; 5 mg), levofloxacin (third generation; 5 mg), aztreonam (30 mg), polymyxin B (10 mg) and colistin (10 mg). MIC_90_s of imipenem, meropenem, ciprofloxacin, chloramphenicol, ceftriaxone, erythromycin and tetracycline were determined by broth microdilution using 2-fold dilution in the range 2–512 mg/L under incubation at 37 °C.

### 2.3. DNA extraction and whole genome sequencing for *P. aeruginosa* DMC30b

The genomic DNA from *P. aeruginosa* DMC30b was extracted from overnight culture by boiled DNA extraction method [26] using commercial DNA extraction kit, QIAamp DNA Mini Kit (QIAGEN, Hilden, Germany). The quality and quantity of the extracted DNA were measured using a NanoDrop ND-2000 spectrophotometer (Thermo Fisher Scientific, Waltham, MA 02451, USA). DNA extracts with A260/280 and A260/230 ratios of ∼ 1.80 and 2.00 to 2.20, respectively, were considered as high-purity DNA sample. Finally, the harvested DNA was visualized on 1% (w/v) agarose gel, and sent for DNA sequencing based on their high purity and adequate concentration.

The whole genome sequencing (WGS) was done under Ion Torrent platform using 400 bp read chemistry [27]. The sequence reactions were performed using the BigDye terminator cycle sequencing kit v3.1 (Applied Biosystems), purified using the BigDye XTerminator purification kit (Applied Biosystems), and then loaded onto a SeqStudio genetic analyzer capillary sequencer (Applied Biosystems) according to the manufacturer’s instructions.

### 2.4. Genome analysis of *P. aeruginosa* strain DMC30b

The Ion Torrent platform generated FASTQ reads quality was initially assessed by the FastQC tool [28] followed by trimming of low-quality sequences using the Trimmomatic [29], while the quality cut off value was Phred-20 [30, 31]. Trimmed reads were assembled de novo using SPAdes v. 3.12 1 [32] in an integrated Genome Analysis Platform (IonGAP) for Ion Torrent sequence data (http://iongap.hpc.iter.es/iongap/newproject). Coverage of the genomes ranged from 76.76x. The circular visualization of the *P. aeruginosa* DMC30b genome was performed using CGViewer [33]. *P. aeruginosa* DMC30b genome was compared with two reference strains; *P. aeruginosa* LESB58 (NCBI sequence accession number NC_011770.1) [34], and *P. aeruginosa* K34-7 (NCBI sequence accession number NZ_CP029707.1) [35] genomes. KmerFinder 3.1 [36] and PathogenFinder 1.1 [37] opensource tools were utilized to identify the species (i.e., *P. aeruginosa*) and pathogenicity of the isolate (DMC30b).

The annotated WGS data were used for sequence typing, antibiotic resistance genes (ARGs) prediction, virulence factor genes (VFGs) profiling, plasmids identification, and metabolic functional analysis. PROKKAAnnotation v.3.2.1 [38] was used to predict the function and identification of assembled sequences against nucleotide and protein sequence database. BUSCO (Benchmarking Universal Single-Copy Orthologs) v.4.1.2 with ‘bacteria_odb10’ data set was used to measure the completeness of the genome [10]. The genetic relatedness of the predicted sequence types, core genome MLST analysis (cgMLST) and bacterial source tracking of the *P. aeruginosa* DMC30b and all of the 166 ST244 *P. aeruginosa* isolates currently deposited in NCBI GenBank database were performed using BacWGSTdb 2.0 server [39]. The Comprehensive Antibiotic Resistance Database (CARD) [40], and VirulenceFinder [41, 42] databases were used to detect ARGs and VFGs, respectively. The CARD (https://card.mcmaster.ca) integrated within AMR++ pipeline identified the respective genes or protein families coding for the ARGs markers in the *P. aeruginosa* DMC30b strain [40]. We utilized SnapGene Viewer web tool (https://www.snapgene.com/snapgene-viewer/) to visualize the virulence plasmid of the *P. aeruginosa* DMC30b. Annotation of the genome was also performed with RAST (Rapid Annotation using Subsystems Technology) server (http://rast.nmpdr.org/) to detect the genomic functional features [43] to detect the genomic functional features.

## 3. Results and Discussion

The KmerFinder 3.1 detected the DMC30b isolate as *P. aeruginosa* DMC30b, while the pathogenicity of the isolate was confirmed (0.973 out of 1.00, close to the pick value indicating higher pathogenicity) by PathogenFinder 1.1. The de novo genome assembly revealed that the *P. aeruginosa* DMC30b genome was 6,994,756 bp with a coverage of 76.76x. The draft genome had 4,595,555 reads assigning for G+C content of 65.7% (Table 1). Analysis of RAST annotated genome using CG view produced a circular genome map illustrating the GC content, tRNA, rRNA, CDS, and contigs (Fig. 1). The genome of *P. aeruginosa* DMC30b possessed 474 contigs including N50 contig 56,583 bp, L50 contig 35 bp, and two plasmids. Of the detected contigs, the longest and shortest contig size were 252,681 bp and 202 bp, respectively. Notably, preliminary sequence analysis revealed 44 insertion sequences (ISs), 21 predicated genomic islands (GIs), six prophage-related sequences and two CRISPR arrays. The genome of *P. aeruginosa* DMC30b encodes for 7,036 genes (Table 1). A physical genome map of *P. aeruginosa* DMC30b in comparison to two other reference strain *P. aeruginosa* LESB58 and *P. aeruginosa* K34-7 is shown in Fig. 1. PROKKA annotated genome using PATRIC produced a circular genome map (Fig. 2), which identified various gene features, 6,685 protein coding sequences (CDSs), 67 RNAs including 4 rRNAs (5S = 1, 16S = 1, 23S = 2), 59 tRNAs, 4 ncRNAs, and 285 pseudogenes. The protein features consist of 3,060 hypothetical proteins, 2,379 functional assignments, and 1,421 GO (gene ontology) assignments. Genome completeness analysis with BUSCO showed the presence of 100% complete BUSCOs in the hybrid assembly.

**Table 1.** *Pseudomonas aeruginosa* strain DMC30b genome characteristics.

| Attribute(s) | Value | % of Total<sup>a</sup> |
| --- | --- | --- |
| Genome size (bp) | 6,994,756 | 100.00 |
| DNA coding (bp) | 6,470,148 | 92.50 |
| Number of reads assemblies | 2,550,071 | 100 |
| Genome coverage | 76.76x | 76.76 |
| DNA G + C (bp) | 4,595,555 | 65.7 |
| Total contigs | 474 |  |
| Longest contig size (bp) | 252,681 | 3.61 |
| Shortest contig size (bp) | 202 | 0.002 |
| Contig N50 (bp) | 56,583 | 0.81 |
| Contig L50 (bp) | 35 |  |
| Total genes | 7,036 | 100.00 |
| Coding sequences (CDSs) | 6,969 | 99.04 |
| CDSs (Protein coding) | 6,685 | 95.01 |
| RNA genes | 67 | 100 |
| tRNA genes | 59 | 88 |
| rRNA genes | 4 | 6 |
| ncRNA genes | 4 | 6 |
| Pseudo genes | 285 | 4.05 |
| Genes with function prediction | 7,146 | 100 |
| Genes assigned to SEED subsystems | 1856 | 26 |
| Number of subsystems | 340 |  |
| CRISPR arrays | 2 |  |
| Number of plasmids | 2 |  |

<sup>a</sup>Total based on either the size of the genome in base pairs (bp) or the total number of genes in the annotated genome.

**Fig. 1.**
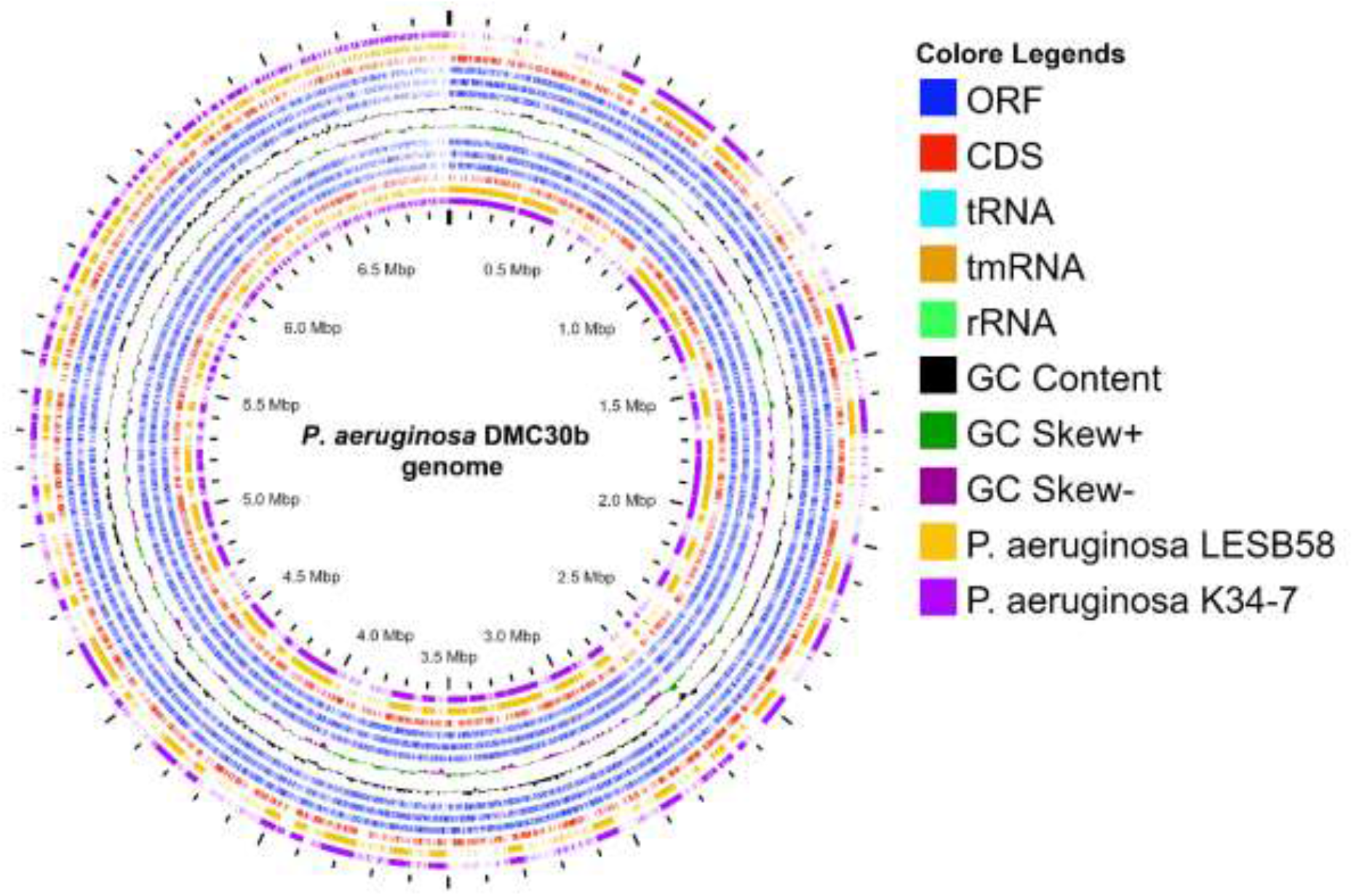
Circular genome representation of *P. aeruginosa* strain DMC30b compared with *P. aeruginosa* LESB58 (GCA_000026645.1) and *P. aeruginosa* K34-7 (NZ_CP029707). The six innermost layers in the graphic portray the genome coordinates (mega base pairs—Mbp, purple), coding sequences (CDS; red), open reading frames (ORF, blue), GC content (zigzag black line) and GC skew (green + /deep purple − zigzag) of the *P. aeruginosa* strain DMC30b genome. The other colored rings, from the outermost to innermost, depict the nucleotide BLAST alignment of *P. aeruginosa* K34-7 (purple) followed by *P. aeruginosa* LESB58 (yellow) representing the positions covered by G+C content (Black), G+C positive skew (green), and G+C negative skew (deep purple). Image created using CGview Server.

**Fig. 2.**
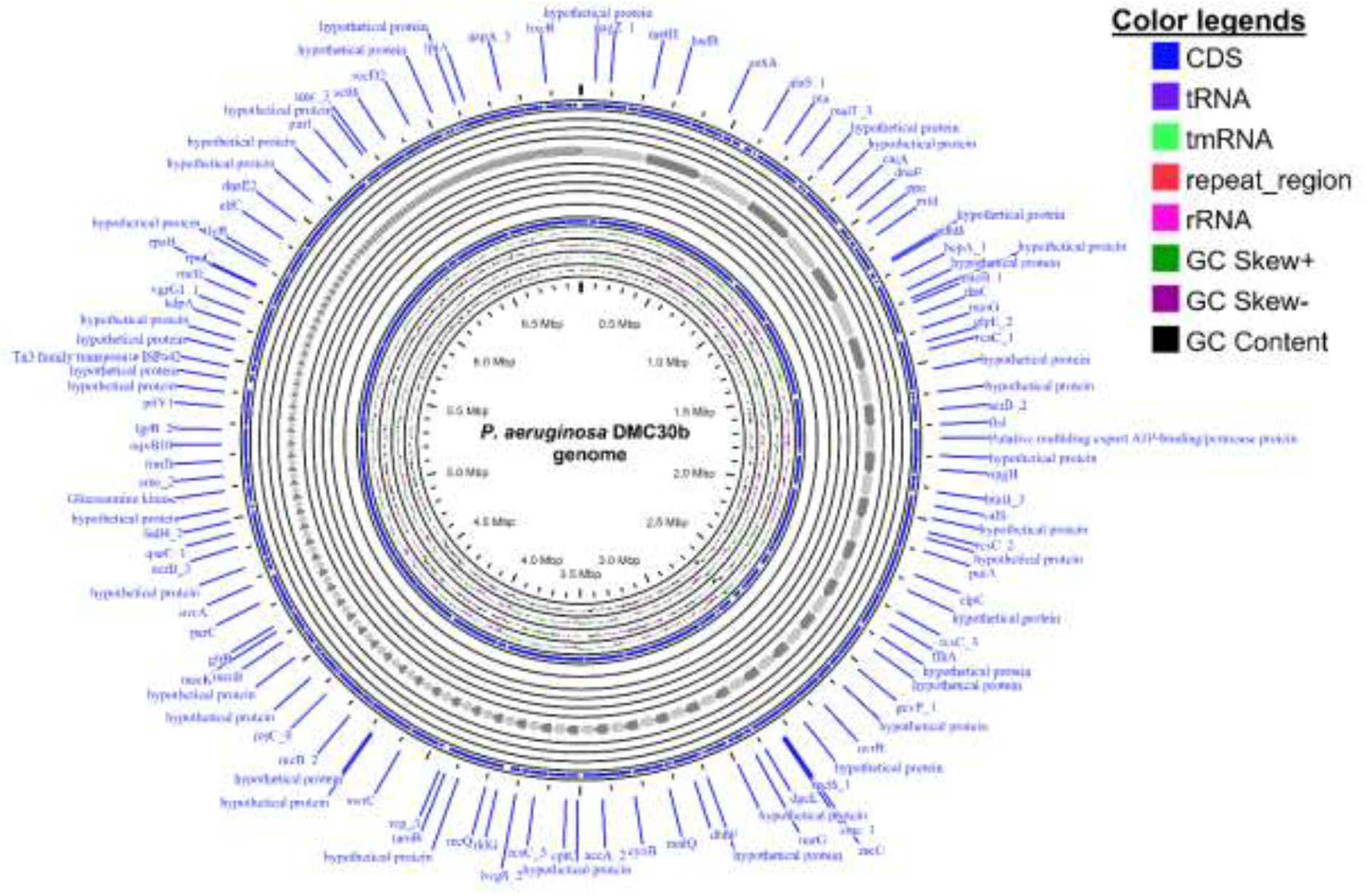
Circular representation of genome *P. aeruginosa* strain DMC30b using CGView Server (http://cgview.ca). The contents are arranged in feature rings (starting with outermost ring): the outermost first ring shows the CDS (coding sequences) with Prokka annotation (both strands combined); tRNA, tmRNA, repeat region and rRNA are indicated; the third ring displays the G+C content; the fourth ring shows the G/C skew information in the (+) strand (green color) and (–) strand (dark pink color).

The diverse and dynamic genetic composition of *P. aeruginosa* enables this bacterium to colonise various environments, including humans where it can cause opportunistic infections [44]. The genomic features of the *P. aeruginosa* DMC30b genome described in this study is corroborated with the genome composition of different *P. aeruginosa* strains reported previously from other countries [34, 35, 45]. For instance, the complete genome of *P. aeruginosa* DMC30b showed 100% identity with two previously reported human originated strains; *P. aeruginosa* LESB58 and *P. aeruginosa* K34-7. The prevalence of MDR or extensively drug resistant strains of *P. aeruginosa* reduces treatment options, significantly increasing morbidity rates. *P. aeruginosa* LESB58, a so-called “Liverpool epidemic strain,” was found to be highly transmissible among cystic fibrosis (CF)-patients and displayed the potential to cause severe infections even in non-CF human hosts [45, 46]. *P. aeruginosa* K34-7 belongs to sequence type 233 (ST233) and is an XDR, carbapenem-resistant clinical isolate expressing the Verona integron-encoded metallo-β-lactamase (VIM-2) [35].

The *P. aeruginosa* DMC30b belongs to ST244 according to Achtman’s MLST scheme for *P. aeruginosa*. To determine the genomic epidemiological characteristics of *P. aeruginosa* strain DMC30b in a global context, the phylogenetic relationships between *P. aeruginosa* DMC30b and a total of 166 other ST244 *P. aeruginosa* strains currently deposited in the NCBI GenBank database were analysed (Fig. 3). cgMLST analyses revealed that the closest relatives of *P. aeruginosa* DMC30b genome were one carbapenem-resistant ST244 *P. aeruginosa* clone in burn patients in Yunnan province, China [17], and another ST244 strain, *P. aeruginosa* MRSN 17623, clinical isolate carrying the blaVIM-6 gene recovered from human in the USA (https://www.pseudomonas.com/strain/show/2623) which differed by 285 cgMLST loci (Fig. 3). For the first time, ST244 is reported emerging in burn patients which exhibited a higher level of resistance to several antibiotics. The obtained results highlight the need to surveillance MDR-resistant isolates in burn patients [17, 47]. Epidemiological source tracking revealed that the *P. aeruginosa* DMC30b strain has close evolutionary relationship with many strains of *P. aeruginosa* originating from a wide arsenal of clinical samples including enteritis, gastroenteritis (diarrheal diseases), urinary tract infections, and both community- and hospital-acquired bacteraemia in humans and animals (e.g., dog). Therefore, *P. aeruginosa* DMC30b may represent a zoonotic threat either by causing disease in human hosts or animal hosts via horizontal gene transfer of plasmid-linked ARGs and/or VFGs to commensal strains.

**Fig. 3.**
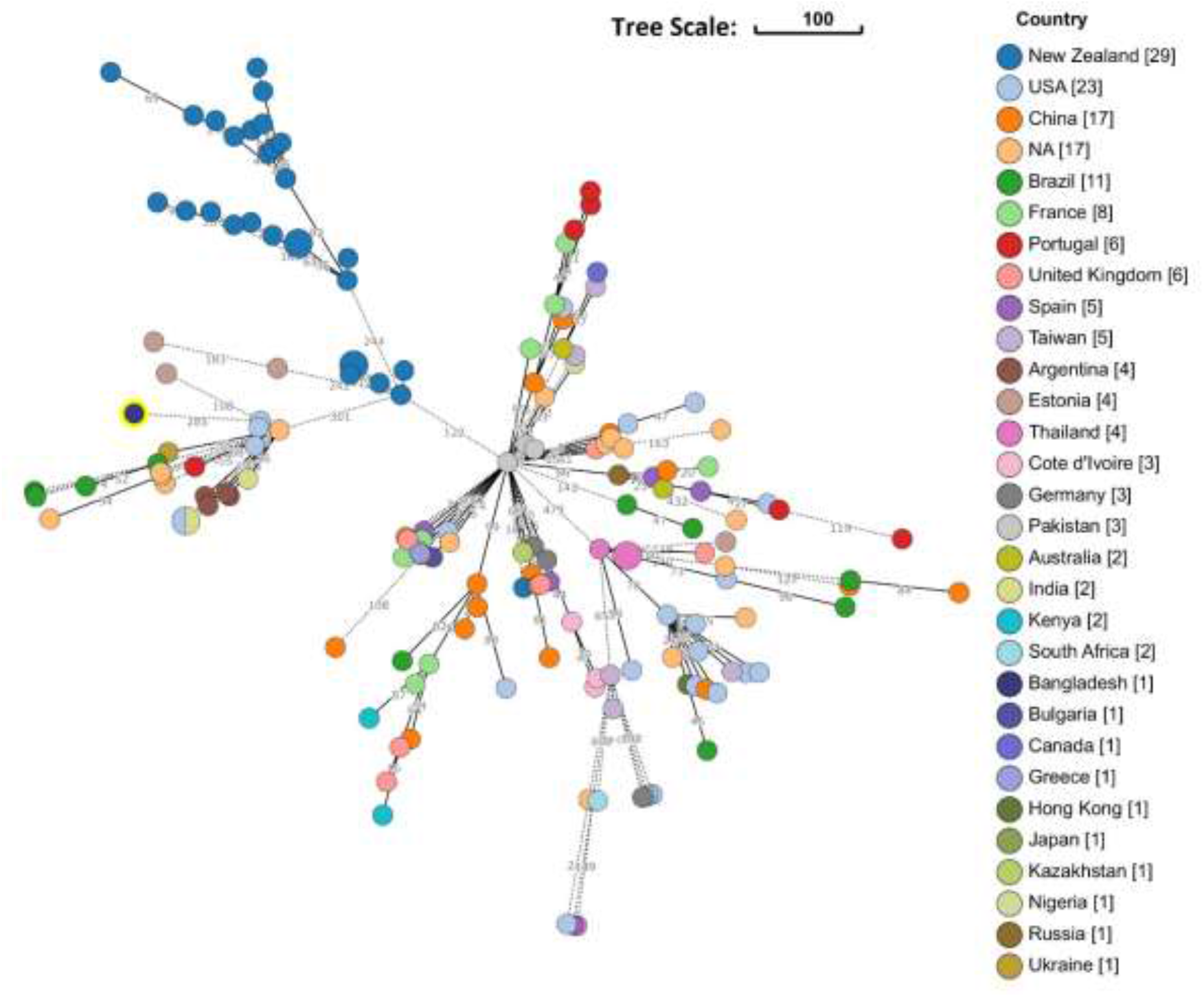
Phylogenetic relationship between *P. aeruginosa* strain DMC30b and all of the 166 *P. aeruginosa* strains belonging to ST244 currently available in the NCBI GenBank database by core genome multilocus sequence typing (cgMLST) analysis.

The genome of the *P. aeruginosa* DMC30b possessed two plasmids namely IncP-6 plasmid p10265-KPC and ColRNAI_pkOIISD1 (Fig. 4). Of them, the IncP-6 plasmid p10265-KPC plasmid was 78,007 bp in size with an average G + C content of 65.7% and 97.59% identity (Fig. 4A) whereas ColRNAI_pkOIISD1 was 9,359 bp in size and showed an identity of 91.35% (Fig. 4B). The IncP-6 plasmid p10265-KPC harboured numerous (> 100) predicted open-reading frames (ORFs) and seven restriction sites for different enzymes with coding sequences (CDSs) (Fig. 4A). BLAST comparison showed that IncP-6 plasmid p10265-KPC of *P. aeruginosa* DMC30b displayed 100% query coverage and 99.60% nucleotide similarity with virulent plasmid p10265-KPC (virulence plasmid of *P. aeruginosa*) isolated in January 2016 from human, China (accession number: KU578314). The plasmid IncP-6 plasmid p10265-KPC is a novel IncP-6 resistance plasmid in the genome of *P. aeruginosa* DMC30b, and carries the IncP-6-type replication, partition and mobilization systems. These results corroborated with previously published results of *P. aeruginosa* strain 10265, recovered from a patient with pneumonia in a public hospital in China [16].

**Fig. 4.**
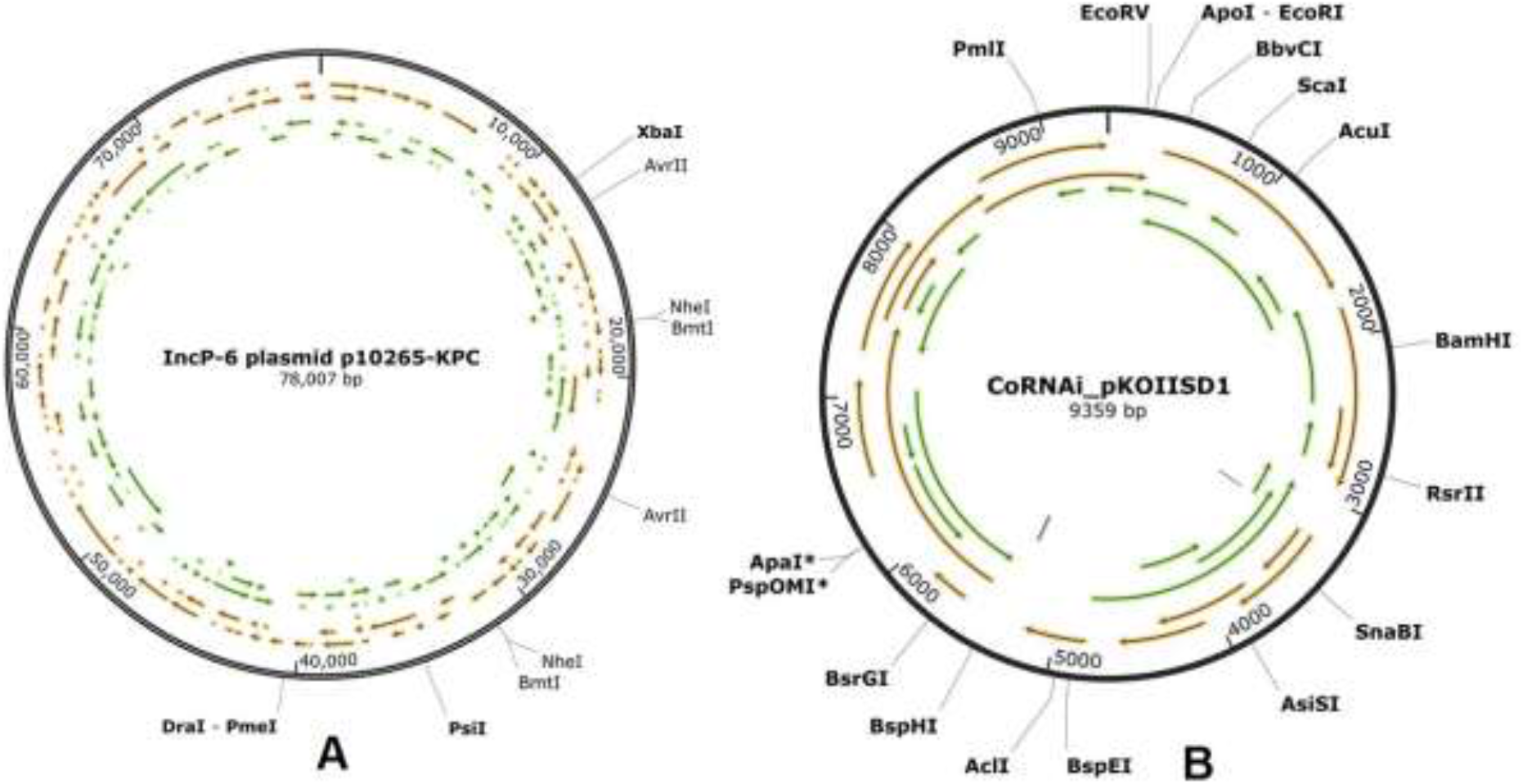
Genetic information surrounding on IncP-6 plasmid p10265-KPC and ColRNAI_pkOIISD1 plasmid of the *P. aeruginosa* DMC30b strain. The arrows denote the open-reading frames (ORFs) and restriction sites for different enzymes with coding sequences (CDSs).

*P. aeruginosa* DMC30b was resistant to 18 antibiotics, with imipenem and meropenem MIC_90_s of >512 mg/L. The *P. aeruginosa* DMC30b was also positive for blaOXA-48 gene. The resistomes of *P. aeruginosa* DMC30b genome consisted of 35 genes (ARGs) conferring for resistance to aminoglycosides (aadA1 and aph(3’)-IIb), four classes of β-lactams (OXA-10, OXA-396, OXA-847, blaPAO, blaPME-1, TEM-116), phenicol (catB7), quinolones (crpP), fluroquinolones (gyrA), fosfomycin (fosA), sulphonamide (sul1), tetracycline (tetD, tetG), bicyclomycin (bcr1), polymyxin (arnA), and colistin (cprR) (Fig. 5). The efflux pump conferring resistance to multiple antibiotics was found as the predominating resistance class in the genome of *P. aeruginosa* DMC30b. For instance, MexAB-OprM, MexCD-OprJ, MexEST-OprN, MexPQ-OpmE, MexIH-OpmD are the leading efflux pump systems conferring resistance to *P. aeruginosa* DMC30b (Fig. 5).

**Fig. 5.**
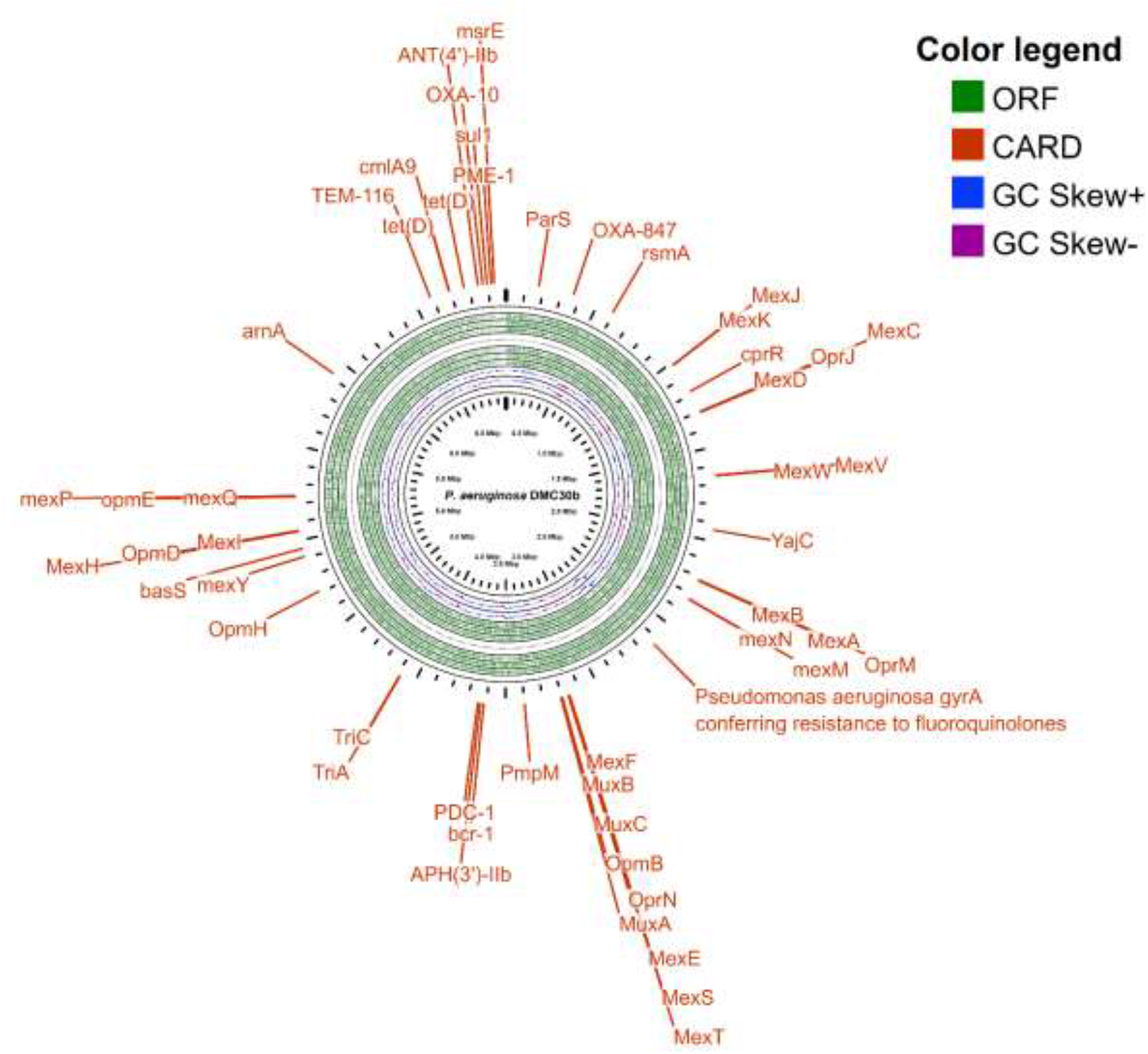
Circular representation of genome *P. aeruginosa* strain DMC30b using CGView Server (http://cgview.ca). The green layer represents the ORF (open reading frame) while GC skew + and GC skew – are depicted by the blue and purple layers, respectively. The outermost layer (black colour coordinates) of the circular graphic shows the antimicrobial resistances genes and related pathways (deep orange) found in the genome through CARD (Comprehensive Antibiotic Resistance Database).

Therefore, according to this study the existence of resistomes (ARGs) and related resistance mechanisms in *P. aeruginosa* DMC30b, along with four classes of beta-lactam genes and efflux pump systems, may contribute to its extensive resistance to almost all antibiotics used for treatment purposes and can cause its emergence as a pandrug-resistant bacterium in Bangladesh. *P. aeruginosa* DMC30b is a particularly pernicious pathogen as it possesses several innate defense mechanisms against antibiotics. These results are in line with our previous findings of *P. aeruginosa* DMC-27b, where the co-existence and chromosomal inheritance of all four b-lactamase classes and acquisition of multiple resistance determinants were reported [8]. This bacterium can readily acquire genetically encoded resistance determinants from other pathogens, further increasing resistance [10, 48]. Clinically significant *P. aeruginosa* strains express resistance to various antimicrobial agents including β-lactam antibiotics through these efflux pump systems [49]. The intrinsic and acquired resistance mechanisms include tripartite efflux systems such as MexAB-OprM, MexCD-OprJ, MexEF-OprN and MexXY/OprM [49, 50] leading to extrusion of antimicrobial agents and other xenobiotics from the cell interior. MexAB-OprM plays a role in the intrinsic resistance of *P. aeruginosa* to most β-lactams, quinolones and many other structurally unrelated antimicrobial agents [49, 50]. Likewise, MexEF-OprN is capable of extruding quinolones [51], and MexXY-OprM expels aminoglycosides and cephalosporin [49, 52] from bacterial cell interior. Furthermore, *P. aeruginosa* achieves high-level (MIC > 1 mg/ml) triclosan resistance either by constitutive expression of triABC, an efflux pump of the resistance nodulation cell division (RND) family, or expression of MexCD-OprJ, MexEF-OprN, and MexJK-OpmH in regulatory mutants, supporting the notion that efflux is the primary mechanism responsible for this bacterium’s high intrinsic and acquired triclosan resistance [53]. Moreover, a series of previous studies reported that most of the ST244 isolates (85.3 %) exhibited a MDR phenotype, i.e. being resistant to carbapenems, aminoglycosides and fluoroquinolones [17, 47].

The genome of *P. aeruginosa* DMC30b harboured 214 VFGs with > 99.0% nucleotide similarity. Of the detected VFGs, genes associated with type III (T3SS) and type VI secretion systems (T6SS), flagella and type IV pili, pyoverdine biosynthesis proteins, flagellar/twitching motility proteins, *Pseudomonas* protein phosphatases (PppA and PppB), two-component regulatory system, transcriptional regulators (TRs), phenazine-modifying enzyme, LasA protease precursor etc. To establish infections, *P. aeruginosa* employs a broad arsenal of virulence determinants, of which its T3SS and T6SS have been the focus of much recent attention [54, 55]. *P. aeruginosa* utilizes T3SS and T6SS apparatuses to inject effector proteins from the its cytosol to the extracellular environment of the host cells to develop acute infections [55, 56]. Pyoverdine, a siderophore produced by *P. aeruginosa*, is essential for pathogenesis in mammalian infections [57], and could represent a novel drug or vaccine target. One of the principle regulatory mechanisms for *P. aeruginosa*’s virulence is the iron-scavenging siderophore pyoverdine, as it governs in-host acquisition of iron, promotes expression of multiple virulence factors, and is directly toxic [48]. Bacterial twitching motility is a surface-associated movement commonly used by Gram-negative bacteria driven by several associated protein functions [58]. Twitching motility, a pilus-mediated form of bacterial surface movement, is required for *P. aeruginosa* virulence in keratitis [59]. Moreover, twitching activity of *P. aeruginosa* play a role not only in motility but also in cell–cell adhesion, cell-surface adhesion and horizontal gene transfer in the pathogenesis of different diseases [60]. The protein phosphatases (PppA and PppB) regulate the basic cellular processes and virulence in *P. aeruginosa* [61]. It has been reported that efflux pumps regulated by two-component systems in several pathogens, including *P. aeruginosa*, provide multidrug resistance, which may limit the treatment options [42, 62]. *P. aeruginosa* encodes a large set of TRs that modulate and manage cellular metabolism to survive in variable environmental conditions including that of the human body, and control the expression of quorum sensing and protein secretion systems [55]. To date, several virulence-related TFs of *P. aeruginosa* have been studied individually, however, little is known about the crosstalk between strictly virulence-related TFs in this bacterial species. Phenazines can modify cellular redox states, regulate patterns of gene expression, enhance survival, contribute to competitiveness, biofilm formation, and virulence in the opportunistic pathogen *P. aeruginosa* [63]. LasA is staphylolytic proteinase which is secreted by the opportunistic pathogen *P. aeruginosa*. LasA protease play a major role in the colonization of the bacteria during bacterial keratitis by preventing other bacteria from colonization to the cornea, for example in the mixed infection with *S. aureus* the enzyme eradicate the bacteria by lysis it and finally eliminate the competitive bacteria for *P. aeruginosa* [64, 65].

The RAST server-based annotation of the *P. aeruginosa* DMC30b genome resulted in a total of 340 subsystems with 26% subsystem coverage (Fig. 6). The SEED functional category distribution of the genes assigned to different subsystems indicated the highest genes encoding for metabolism of amino acids and derivatives (506 genes), followed by the metabolism of carbohydrates (287 genes), protein metabolism (223 genes), cofactors, vitamins, prosthetic group, and pigments (202 genes), metabolism of aromatic compounds (123 genes), DNA metabolism (103 genes) and fatty acids, lipids, and isoprenoids (97 genes). RAST based annotation classified 182 genes in the subsystem category ‘membrane transport’, 124 genes in subsystem category ‘respiration’ and 108 genes in subsystem category ‘stress response’ (Fig. 6). Further analysis of the genes in the stress response subsystem category revealed the presence of genes associated with oxidative stress (∼ 60%), osmotic stress (12.0%) and periplasmic stress (∼ 6%). The *P. aeruginosa* DMC30b genome also encodes for 69 genes for virulence, disease and defense in the subsystem analysis (Fig. 6), of which 52 (75.36%) genes were associated with resistance to antibiotics and toxic compounds, 14 (20.29%) genes for invasion and intracellular resistance, and 3 (4.34%) genes for antibacterial peptides.

**Fig. 6.**
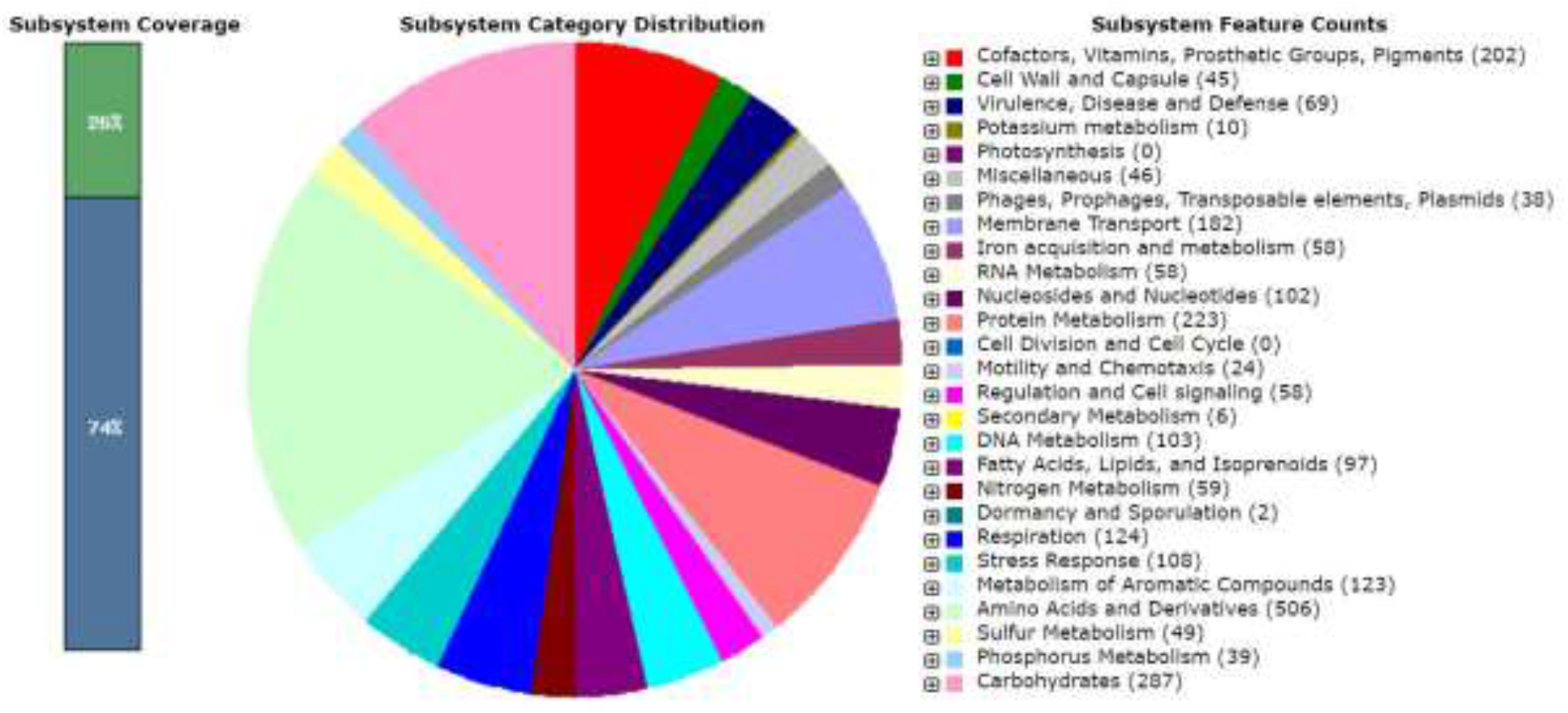
An overview of the subsystem categories assigned to the genes predicted in the genome of *P. aeruginosa* strain DMC30b. The whole-genome sequence of the strain DMC30b was annotated using the RAST server.

We found genes and proteins from the complete genome analysis of *P. aeruginosa* DMC30b attributable to different subsystem metabolic functions which corroborated with several previous studies [17, 54, 66]. Our analysis provides an important expansion of metabolic functional genes and/or pathways integral to the development of infection by *P. aeruginosa*. The existence of multiple *P. aeruginosa* virulence factors and their direct metabolic regulators are linked to pathogenicity. Estimating the percentage of the core metabolism genes participating in the virulence of *P. aeruginosa* DMC30b, we found that genes associated with metabolism are correlated with pathogenicity. Virulence-linked pathways in opportunistic pathogens are putative therapeutic targets that may be associated with less potential for resistance than targets in growth-essential pathways [17, 66]. However, efficacy of virulence-linked targets may be affected by the contribution of virulence-related genes to metabolism [66]. Furthermore, given its capacity to metabolize a wide variety of substrates, it is also possible that *P. aeruginosa* DMC30b possesses greater potential for enzymatic modification, virulence and MDR mechanisms. Therefore, the metabolic diversity, transport capabilities and regulatory adaptability that enable *P. aeruginosa* DMC30b to thrive and compete with other microorganisms probably all contribute to its high intrinsic resistance to antibiotics. Knowledge of the complete genome sequence and encoded processes (ARGs, VFGs and subsystems) provides a wealth of information for the discovery and exploitation of new antibiotic targets, and hope for the development of more effective strategies to treat the life-threatening opportunistic infections caused by *P. aeruginosa* in humans.

In summary, this is the first WGS-based study on MDR *P. aeruginosa* DMC30b isolated from septic wound swab of a severely affected burn patient in Bangladesh. In this study, we report the genomic characteristics of a MDR *P. aeruginosa* DMC30b carrying different ARGs, VFGs and genomic functional potentials which may help to elucidate the dissemination mechanisms of resistance and virulent genes among bacteria, animals, and humans. These data can facilitate unravelling the genomic features, antimicrobial resistance mechanisms and epidemiological characteristics of this bacterial pathogen in the future. Further studies are required to have a wider understanding of genetic features of *P. aeruginosa* isolates circulating within communities in Bangladesh. This could ultimately bring about informed government policy and adequate community awareness toward curbing the transmission and continuous emergence of such dangerous pathogens.

## 4. Data availability

The WGS data of the *P. aeruginosa* DMC30b is deposited at DDBJ/ENA/GenBank under accession number JAMQYG000000000 (BioSample SAMN28906490), and the assembly reports of the genome are also available at GenBank (https://ncbi.nlm.nih.gov/nuccore/JAMQYG000000000). The version described in this paper is version JAMQYG000000000.1. The Ion Torrent FASTQ reads are available in the National Center for Biotechnology Information (NCBI) under BioProject accession number PRJNA846956.

## Acknowledgments

The authors would like extend their gratitude to TWAS (The World Academy of Science) for funding research grants to complete the study. We, also would like to thank Sumaiya Sharmin (Lecturer, Primeasia University, Bangladesh) for her assistance in collecting the study isolates from Dhaka Medical College (DMC) Hospital, Bangladesh. Our sincere gratitude to Dr. Zhi Ruan (Sir Run Run Shaw Hospital, Zhejiang University School of Medicine, Hangzhou 310016, China) for his technical support in BacWGSTdb 2.0 analysis of the genome.

